# Prefrontal Cortex Regulates Chronic Stress-Induced Cardiovascular Susceptibility

**DOI:** 10.1101/675835

**Authors:** Derek Schaeuble, Amy E.B. Packard, Jessica M. McKlveen, Rachel Morano, Sarah Fourman, Brittany L. Smith, Jessie R. Scheimann, Ben A. Packard, Steven P. Wilson, Jeanne James, David Y. Hui, Yvonne M. Ulrich-Lai, James P. Herman, Brent Myers

## Abstract

The medial prefrontal cortex (mPFC) is necessary for appropriate appraisal of stressful information, as well as coordinating visceral and behavioral processes. However, prolonged stress impairs mPFC function and prefrontal-dependent behaviors. Additionally, chronic stress induces sympathetic predominance, contributing to health detriments associated with autonomic imbalance. Previous studies identified a subregion of rodent prefrontal cortex, infralimbic cortex (IL), as a key regulator of neuroendocrine-autonomic integration after chronic stress, suggesting that IL output may prevent chronic stress-induced autonomic imbalance. In the current study, we tested the hypothesis that the IL regulates hemodynamic, vascular, and cardiac responses to chronic stress. To address this hypothesis, a viral-packaged siRNA construct was used to knockdown vesicular glutamate transporter 1 (vGluT1) and reduce glutamate packaging and release from IL projection neurons. Male rats were injected with a vGluT1 siRNA-expressing construct or GFP control into the IL and then remained as unstressed controls or were exposed to chronic variable stress (CVS). IL vGluT1 knockdown increased heart rate and mean arterial pressure (MAP) reactivity, while CVS increased chronic MAP only in siRNA-treated rats. In a separate cohort, CVS and vGluT1 knockdown interacted to impair both endothelial-dependent and endothelial-independent vasoreactivity *ex vivo*. Furthermore, vGluT1 knockdown and CVS increased histological markers of fibrosis and hypertrophy. Thus, knockdown of glutamate release from IL projection neurons indicates that these cells are necessary to prevent the enhanced physiological responses to stress that promote susceptibility to cardiovascular pathophysiology. Ultimately, these findings provide evidence for a neurobiological mechanism mediating the relationship between stress and poor cardiovascular health outcomes.

**Clinical perspective:** What is new?

- Knockdown of glutamate release from infralimbic cortex increases heart rate and arterial pressure reactivity
- Decreased infralimbic glutamate output leads to vascular dysfunction after chronic stress
- These functional changes associate with histological indicators of cardiac hypertrophy, as well as vascular hypertrophy and fibrosis

What are the clinical implications?

- These studies provide a neurobiological mechanism that may account for the link between long-term stress and increased cardiovascular disease risk

## Introduction

Stress, a real or perceived threat to homeostasis or well-being, elicits behavioral and physiological responses to promote organismal adaptation^1, 2^. However, prolonged stress exposure has deleterious effects on health, increasing susceptibility to cardiovascular, psychiatric, and metabolic disorders^3–6^. In fact, chronic psychosocial stress predicts the incidence of cardiovascular disease, cardiac-related morbidity and mortality, and doubles the risk of myocardial infarction^7, 8^. Exaggerated heart rate (HR) reactivity to acute stress also predicts poor cardiovascular outcomes, including hypertension, ventricular hypertrophy, and atherosclerosis^9^. Although the biological mechanisms mediating the relationship between stress and cardiovascular health are not completely understood, adverse outcomes likely result from prolonged exposure to neural and endocrine stress mediators.

The initial appraisal of psychological stressors largely occurs in the limbic system, a network of interconnected structures spanning the forebrain. The medial prefrontal cortex (mPFC) is a key limbic cortical structure mediating stress appraisal, emotion, and cognition^10–13^. Moreover, activity within a specific region of the ventral mPFC, the subgenual cingulate cortex (BA25), associates with sadness in healthy controls^14^, as well as pathological depression in treatment-resistant patients^15^. Recent human neuroimaging studies have also identified the ventral mPFC as a component of the central autonomic network that responds to and coordinates visceral functions, including stress-evoked blood pressure reactivity^16–19^. The rodent homolog of BA25 is infralimbic cortex (IL)^20–22^. This subregion of ventral mPFC provides inputs to stress-integrative nuclei, including the posterior hypothalamus and brainstem autonomic centers^23–26^. Our previous studies reduced glutamate outflow from the IL in rats undergoing chronic variable stress (CVS) and found hyperactivation of the hypothalamic-pituitary-adrenal axis (HPA) axis^27, 28^. As glutamate release from IL projections is a key regulator of acute and chronic neuroendocrine reactivity, we hypothesized that IL output may prevent chronic stress-induced autonomic imbalance and associated cardiovascular susceptibility.

To address this hypothesis, a lentiviral-packaged small interfering RNA (siRNA) targeting vesicular glutamate transporter 1 (vGluT1) was injected in the IL. This approach selectively reduces vGluT1 expression in IL glutamate neurons, preventing the packaging and release of glutamate from presynaptic terminals^28–30^. Animals were then exposed to CVS to examine interactions between chronic stress and hypo-functionality of ventral mPFC in terms of cardiovascular reactivity, arterial function, and remodeling of the vasculature and myocardium. These studies identified the necessity of the IL for reducing hemodynamic, vascular, and cardiac consequences of prolonged stress. Additionally, this work points toward a neurobiological mechanism mediating the relationship between stress and cardiovascular health.

## Methods

The data that support the findings of this study are available from the corresponding author upon reasonable request.

### Animals

Adult male Sprague-Dawley rats were obtained from Harlan (Indianapolis, IN) with weights ranging from 250-300 g. Rats were housed individually in shoebox cages in a temperature- and humidity-controlled room with a 12-hour light-dark cycle (lights on at 0600h, off at 1800h) and food and water *ad libitum*. All procedures and protocols were approved by the University of Cincinnati Institutional Animal Care and Use Committee (protocol: 04-08-03-01) and complied with the National Institutes of Health Guidelines for the Care and Use of Laboratory Animals. The cumulative sequence of procedures employed in the current experiments received veterinary consultation and all animals had daily welfare assessments by veterinary and/or animal medical service staff. Signs of poor health and/or weight loss ≥ 20% of pre-surgical weight were *a priori* exclusion criteria. These criteria were not met by any animals in the current experiments.

#### Experiment 1

##### Design

For experiment 1, 32 rats (n = 8/group) were injected with either a lentiviral-packaged construct coding for vGluT1 siRNA or GFP as a control. After instrumentation with radiotelemetry devices, half of the animals were exposed to 14 days of CVS with the rest of the animals remaining as No CVS controls. All treatment assignments were randomized. Home cage cardiovascular parameters were continuously monitored in all rats throughout the 14-day period of CVS. On the morning of day 15, all rats were subject to an acute novel restraint to examine hemodynamic stress reactivity.

##### Viral construct

A lentivirus transfer vector, based on a third-generation, self-inactivating transfer vector was constructed as previously described^25, 28^. Briefly, a 363-bp piece of DNA from the rat vGluT1 complementary DNA was synthesized that included 151 bp of the 3’ coding region and 212 bp of the 3’ untranslated region. This corresponds to nucleotides 1656 to 2018 of GenBank accession no. NM_053859. This is a region of low homology with vGluT2 and vGluT3 and avoids all the putative transmembrane domains of the transporter. The fragment was cloned in antisense orientation into a lentivirus transfer vector that expressed an enhanced green fluorescent protein (GFP) reporter. This vector uses the phosphoglycerate kinase-1 promoter, which expresses well in rat brain and is primarily neuronal^28, 31^. A control virus was constructed similarly, using a transfer vector with the phosphoglycerate kinase-1 promoter driving expression of enhanced GFP alone.

##### Stereotaxic surgery

Animals were anesthetized (90 mg/kg ketamine and 10 mg/kg xylazine, intraperitoneal) followed by analgesic (2 mg/kg butorphanol, subcutaneous) and antibiotic (5 mg/kg gentamicin, intramuscular) administration. Rats received bilateral 1 μL microinjections (5 x 10^6^ tu/μL titer) into the IL (2.9 mm anterior to bregma, 0.6 mm lateral to midline, and 4.2 mm ventral from dura), as described previously^23, 28, 32^, of either the vGluT1 siRNA virus or GFP control. All injections were carried out with a 25-gauge, 2-μL microsyringe (Hamilton, Reno, NV) using a microinjection unit (Kopf, Tujunga, CA) at a rate of 5 minutes/μL. To reduce tissue damage and allow diffusion, the needle was left in place for 5 minutes before and after injections. Animals recovered for 6 weeks before commencing experiments, corresponding to timeframes previously used for similar lentiviral systems^28, 33^.

##### Telemetry

Four weeks after stereotaxic surgery, rats were implanted with radiotelemetry transmitters (PA-C40; Data Sciences International, St. Paul, MN) as previously described^34, 35^. Briefly, animals were anesthetized with inhaled isoflurane anesthesia (1-5%). The descending aorta was exposed via an abdominal incision, allowing implantation of a catheter extending from the transmitter. The catheter was secured with tissue adhesive (Vetbond; 3M Animal Care Products, St. Paul, MN) and a cellulose patch. The transmitter body was then sutured to the abdominal musculature, followed by suturing of the abdominal musculature and closure of the skin with wound clips. Rats recovered for 2 weeks before wound clips were removed and experiments began.

##### Chronic variable stress

CVS was comprised of twice daily (AM and PM) repeated and unpredictable stressors presented in a randomized manner, including exposure to a brightly-lit open field (1 m^2^, 5 minutes), cold room (4°C, 1 hour), forced swim (23° to 27°C, 10 minutes), brightly-lit elevated platform (0.5 m, 5 minutes), shaker stress (100 rpm, 1 hour), and hypoxia (8% oxygen, 30 minutes). Additionally, overnight stressors were variably included, comprised of social crowding (6-8 rats/cage, 16 hours) and restricted housing (mouse cage, 16 hours). All animals went through the CVS paradigm concurrently. To prevent body weight differences between stress conditions, rats remaining unstressed in the home cage were food restricted in accordance with the reduced food intake induced by CVS^34, 36^. During the 2 weeks of CVS, unstressed animals received 80% of a food allotment prior to lights off and the other 20% after lights on to reduce the potential for fasting^34, 36^. On day 15, all rats were exposed to a novel acute restraint to directly compare the effects of vGluT1 knockdown on cardiovascular responses to acute and chronic stress.

##### Acute stress

The morning after completion of CVS (approximately 16 hours after the last stress exposure), all animals were subjected to a novel 40-minute restraint. Stress response assessment was initiated between 08:00 and 09:00 hours. Animals were placed in well-ventilated Plexiglass restraint tubes with baseline pressure and HR measurements collected in the one-hour period preceding restraint. After restraint, rats were returned to their home cage with pressure and HR recorded for 60 minutes after restraint.

##### Tissue collection

After acute restraint, all animals were euthanized with sodium pentobarbital (≥ 200 mg/kg, intraperitoneal) and transcardially perfused with 0.9% saline followed by 4% phosphate-buffered paraformaldehyde. Brains were postfixed in paraformaldehyde for 24 hours and then stored in 30% sucrose at 4°C. Brains were subsequently sectioned (30 μm-thick coronal sections) and processed for GFP immunohistochemistry to determine microinjection spread.

##### Immunohistochemistry

For single immunolabeling of GFP, tissue sections were washed in 50 mM KPBS and incubated in blocking buffer (50 mM KPBS, 0.1% bovine serum albumin, and 0.2% TritonX-100) for 1 hour at room temperature. Sections were placed in rabbit anti-GFP primary antibody (1:1000 in blocking buffer; Invitrogen, La Jolla, CA) overnight at 4°C. Following incubation, sections were rinsed and placed into Alexa488-conjugated donkey anti-rabbit immunoglobulin G (IgG; 1:500 in blocking buffer; Jackson Immunoresearch, West Grove, PA) for 30 minutes. Sections were rinsed, mounted onto slides, and cover slipped. Dual fluorescent immunolabeling was performed as described previously^28^, with GFP labeled in sequence with vGluT1. vGluT1 was visualized with rabbit anti-vGluT1 primary antibody (1:1000; Synaptic Systems, Goettingen, Germany) followed by Cy3-conjugated donkey anti-rabbit IgG (1:500; Jackson ImmunoResearch, West Grove, PA).

##### Microscopy

For visualization of GFP and vGluT1 co-localization, digital images were captured from a 1-in-12 series with a Zeiss Axio Imager Z2 microscope using optical sectioning (63x objective) to permit co-localization within a given z-plane (0.5-μm thickness). Co-localizations were defined as white fluorescence from overlap between labeled GFP terminals and magenta-colored vGluT1. For each figure, brightness and contrast were enhanced uniformly using Adobe Photoshop (CC 14.2).

##### Data analysis

Data are expressed as mean +/- standard error of the mean (SEM). Quantification was conducted by experimenters blind to conditions. All analyses were conducted using GraphPad Prism (version 7.04) for 2-way ANOVA, R Studio (version 3.4.2) for 3-way repeated measures ANOVA, or Dataquest A.R.T. (version 4.3) for telemetry analysis. Over the course of CVS, activity, HR, arterial pressures, and pulse pressure were sampled in the home cage. Samples were collected during the light phase from 06:00-08:00 for AM measures (prior to the first stressor of the day) and during the dark phase from 19:00-21:00 for the PM period (at least 2 hours after the second stressor of the day). During each 2-hour time period, samples were averaged into 10-minute bins for analysis. Beginning the day before CVS, circadian curves were generated for each parameter over the 15-day period. Data over time (both CVS and acute restraint) were analyzed by 3-way repeated measures analysis of variance (ANOVA); with viral treatment, stress, and time (repeated) as factors. When significant main effects were reported, ANOVA was followed by Tukey post-hoc test to identify specific group differences. Area under the curve (AUC) for the 15 days of CVS or 100 minutes of acute stress was calculated by summing the average values acquired between two time points multiplied by the time elapsed [Σ((Time 1 + Time 2)/2)*Time elapsed]. Statistical significance for cumulative measures was determined by 2-way ANOVA with treatment and stress as factors. ANOVA was followed by Tukey post-hoc tests in the case of significant main effects. Values more than 2 standard deviations from the mean were removed as outliers. No animals were removed from the study as outliers but specific time points for telemetry recordings were identified as outliers based on deviation from the mean. These exclusion criteria were developed *a priori* and applied uniformly. The excluded data points were found to represent non-physiological parameters (e.g. HR < 200 beats/minute). Statistical significance was reported as (p < 0.05) for all tests.

#### Experiment 2

##### Design

Experiment 2 employed a similar design as experiment 1. Male rats (n = 7/group) received IL injections of lentiviral-packaged vGluT1 siRNA or GFP and experienced CVS or remained unstressed. All treatment assignments were randomized. On day 15, thoracic aorta was collected to examine vasoreactivity and histology. Additionally, hearts and brains were collected for histological analyses.

##### Chronic variable stress

For experiment 2, CVS was staggered based on the throughput of vascular function experiments. Seven cohorts (n = 4 animals/cohort, one animal for each treatment group) began the CVS paradigm one day apart so that each cohort of 4 would have tissue collected on successive days. All rats undergoing CVS in experiment 2 received the same stressors in the same sequence. The CVS paradigm was similar to experiment 1, except for the substitution of restraint (30 minutes) for crowding.

##### Tissue collection

The morning of day 15, approximately 16 hours after the last stress exposure, all animals were rapidly anesthetized with isoflurane (5%) and decapitated. Thoracic aorta was collected by dissecting 4 mm of aortic tissue proximal to the diaphragm for vascular function analysis. Additional aortic tissue was collected proximal to the initial sample and post-fixed in paraformaldehyde for histological analysis. Hearts were also collected and post-fixed in paraformaldehyde for histological analysis. Brains were post-fixed and subsequently sectioned and processed for GFP immunohistochemistry to determine microinjection spread as described for experiment 1.

##### Vascular function

Aortic tissue samples were processed for wire myographic vasoreactivity analyses as previously described^37^. Briefly, vessels were kept in warm, oxygenated Krebs’s solution with connective tissue and adipose removed under a dissecting microscope. Aortic rings were then placed on wires in organ baths (Radnoti, Covina, CA), equilibrated, and brought to tension. Vasoconstriction was assessed in response to increasing concentrations of the endothelial-dependent potassium chloride (KCl; 0 to 50 mM) and endothelial-independent phenylephrine (1×10^-6^ to 1×10^-2^ mM). Vessels were then set to 80% of maximum phenylephrine-induced constriction and relaxation was determined in response to endothelial-dependent acetylcholine (1×10^-6^ to 1×10^-2^ mM) and endothelial-independent sodium nitroprusside (SNP; 1×10^-7^ to 3×10^-3^ mM). At the end of the experiment, all vessels were weighed and measured (length, circumference, and area) to verify equal dimensions across all groups.

##### Histology and microscopy

Cardiac and aortic tissue was processed by the Cincinnati Children’s Hospital Medical Center Research Pathology Core. Briefly, aortas were paraffin-embedded, sectioned (5 μm), and stained. Verhoeff-Van Gieson (VVG) was used to quantify elastin in the tunica media and Masson’s Trichrome to quantify collagen in the tunica adventitia as previously described^35, 37^. Paraffin-embedded hearts were oriented for four-chamber view, sectioned (5 μm), and stained with Masson’s Trichrome to visualize collagen and wheat germ agglutinin (WGA) conjugated to Alexa488 to visualize myocyte cell membranes as previously described^37, 38^. Vascular and heart tissue were imaged with a Zeiss AxioObserver microscope using a color camera and 10x objective.

##### Data analysis

Data are expressed as mean ± SEM. Quantification was conducted by experimenters blind to conditions. All analyses were conducted using GraphPad Prism (version 7.04) for 2-way ANOVA, R Studio (version 3.4.2) for 3-way repeated measures ANOVA, or FIJI (version 1.51N) for histological quantification. Vasoreactivity data were analyzed by 3-way repeated measures ANOVA; with viral treatment, stress, and drug concentration (repeated) as factors. When significant main effects were reported, ANOVA was followed by Tukey post-hoc test to identify specific group differences. FIJI (version 1.51N) was used to quantify lumen and tunica media dimensions in VVG-stained aorta, as well as adventitia dimensions in Masson’s Trichrome-stained tissue. FIJI was also used to quantify collagen density in hearts stained with Masson’s Trichrome. For each animal, 6 sections of aorta or heart were quantified and averaged. To determine myocyte surface area, FIJI was used to binarize myocyte images. This technique produced dark cytoplasm and bright membranes in cardiomyocytes^39^. The dark cytoplasm was used to calculate surface areas of the myocytes in the apex and lateral wall of the left ventricle. For each animal, approximately 100 cells were counted from 6 heart sections and averaged. Statistical significance for histological measures was determined by 2-way ANOVA with treatment and stress as factors. ANOVA was followed by Tukey post-hoc tests in the case of significant main effects. Statistical significance was reported as (p < 0.05) for all tests.

## Results

### vGluT1 knockdown

Injections of a lentiviral-packaged construct expressing vGluT1 siRNA were targeted to the IL (Fig. 1A, 1B). Viral injections were largely limited to the deep layers of IL, with minimal spread to the prelimbic cortex (PL). We have previously shown that this approach reduces vGluT1 mRNA specifically in the IL, as well as vGluT1 protein co-localization with GFP-labeled terminals^28^. In the current study, tissue from rats injected with a GFP control construct exhibited substantial co-localization with vGluT1 protein on cortico-cortical axonal processes (Fig. 1C). IL projections transfected with the vGluT1 siRNA construct had reduced co-localization with vGluT1 protein (Fig. 1D).

**Figure 1.**
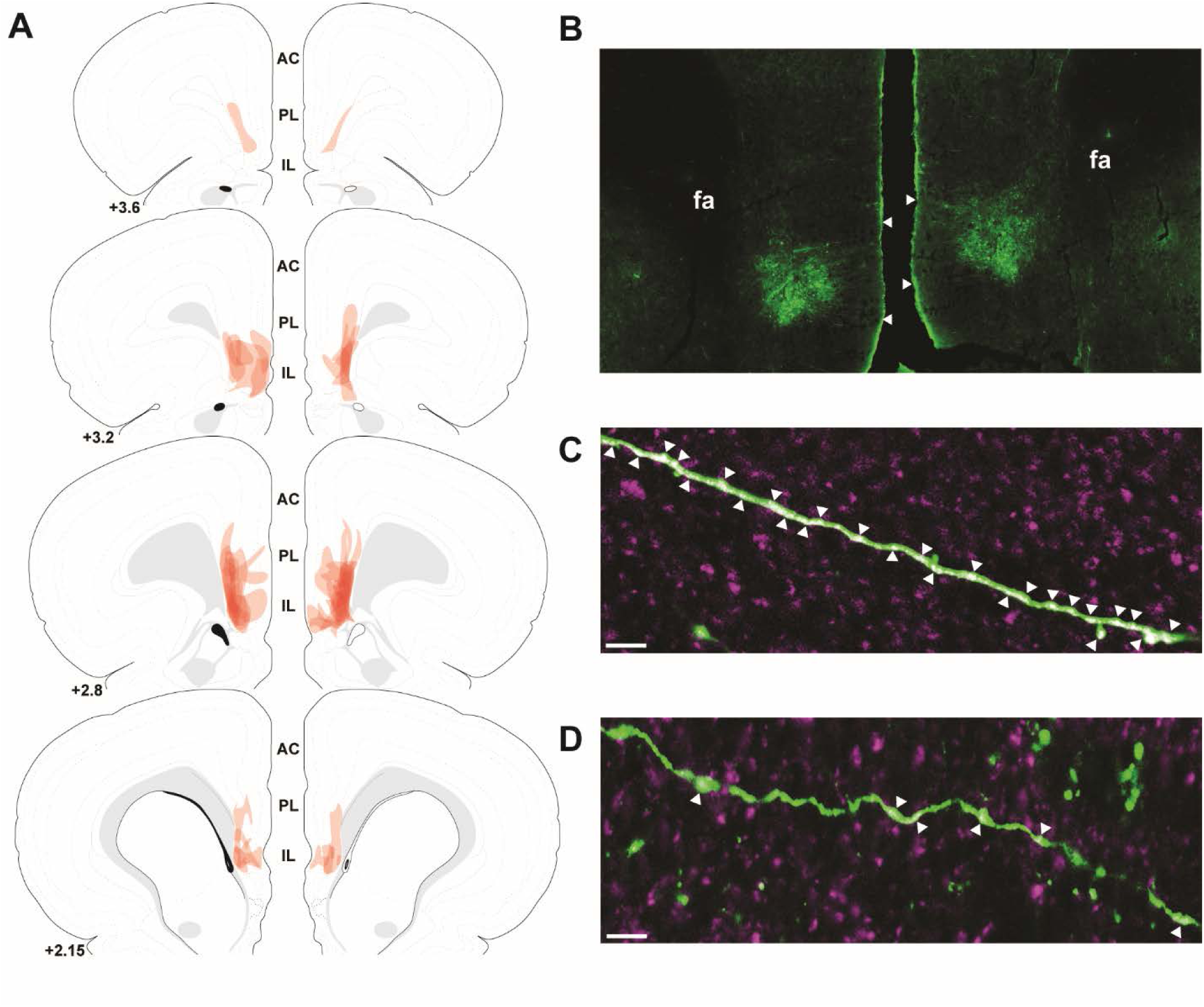
The spread of individual lentiviral injections was traced on photomicrographs and overlaid onto atlas templates from Swanson^74^ to depict the localization of vGluT1 knockdown in experiment 1 (A). Lentiviral injections targeted to the IL with minimal spread to the PL (B). White arrows indicate dorsal and ventral boundaries of the IL. Immunolabeling of GFP (green) and vGluT1 (magenta) indicated a high-degree of co-localization (white arrows) on IL projections in GFP controls (C). Knockdown of vGluT1 with siRNA treatment decreased vGluT1 co-localization with GFP on IL projections (D). Scale bars: (B) 100 µm, (C,D) 10 µm. Numbers indicate distance rostral to bregma in millimeters. AC: anterior cingulate, PL: prelimbic cortex, IL: infralimbic cortex, fa: anterior forceps of the corpus callosum.

### Body weight and food intake

Chronic stress reduces food intake and body weight gain, leading to significant differences in body composition compared to control animals^27, 28, 34, 40^. As this may confound results related to HR and blood pressure reactivity^34^, animals in the No CVS groups for both experiments 1 and 2 received mild food restriction to match body weight with CVS rats (Table 1). In both experiments, there were no significant differences in body weight between groups. However, food restriction in experiment 1 led to food consumption that was significantly greater than the CVS groups [F(1,28) = 31.24, p < 0.0001]. In experiment 2, food restriction significantly decreased food intake compared to CVS groups [F(1,24) = 33.25, p < 0.0001].

**Table 1.**
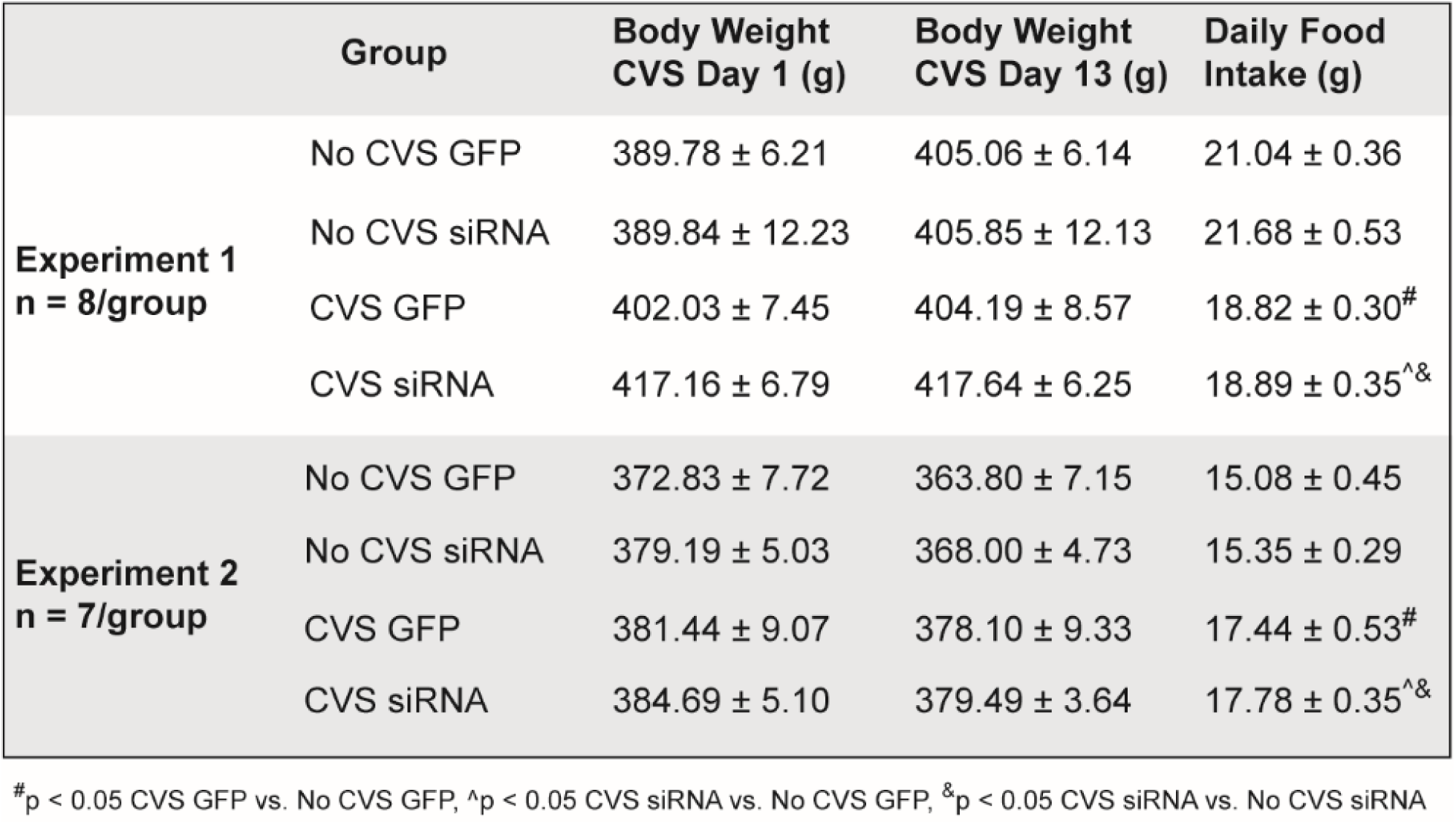
Body weight and food intake throughtout chronic variable stress. Body weight of animals at the beginning and end of CVS for experiments 1 and 2. In both experiments, CVS rats had *ad libitum* access to chow while No CVS animals received mild food restriction to prevent significant differences in body weight between chronically stressed animals and controls.

#### Experiment 1

##### Circadian behavioral activity

Home cage radiotelemetry data were analyzed for 15 days beginning the day before CVS. Throughout CVS, 3-way repeated-measures ANOVA of circadian activity (n = 7-8/group) revealed a main effect of time [F(29, 930) = 21.38, p < 0.0001] and interactions of stress x treatment [F(1, 930) = 4.67, p = 0.031] and stress x time [F(29, 930) = 4.34, p < 0.0001]. Post-hoc analysis indicated that, during the dark period of CVS day 2, rats that were subjected to an overnight crowding stressor exhibited more activity than No CVS controls (Fig. 2A). The CVS GFP rats had elevated activity compared to No CVS GFP controls (p = 0.014); furthermore, CVS siRNA animals were more active than CVS GFP rats (p = 0.007). In addition to overnight social crowding on day 2, rats experienced overnight housing in mouse cages days 6 and 10. Throughout the dark periods of days 6-9, CVS siRNA rats had decreased activity compared to No CVS animals (p ≤ 0.047). Additionally, the CVS GFP rats were less active on the dark periods of days 9 (p = 0.008) and 14 (p = 0.003) compared to No CVS GFP. Also, on the dark period of Day 14, the CVS siRNA group was more active than CVS GFP (p = 0.003). Additional analysis was carried out on cumulative activity counts (No CVS GFP: 801.88 ± 1.04, No CVS siRNA: 819.29 ± 8.08, CVS GFP: 683.66 ±1.38, CVS siRNA: 860.81 ± 4.46). According to 2-way ANOVA, there were significant effects of treatment [F(1,27) = 338.60, p < 0.0001], stress [F(1,27) = 60.38, p < 0.0001], and a treatment x stress interaction [F(1,27) = 261.9, p < 0.0001]. Tukey post-hoc test indicated that total activity over 15 days was lower in CVS GFP rats compared to No CVS GFP (p < 0.0001). In contrast, cumulative activity was increased in CVS siRNA rats compared to all other groups (p < 0.0001).

**Figure 2.**
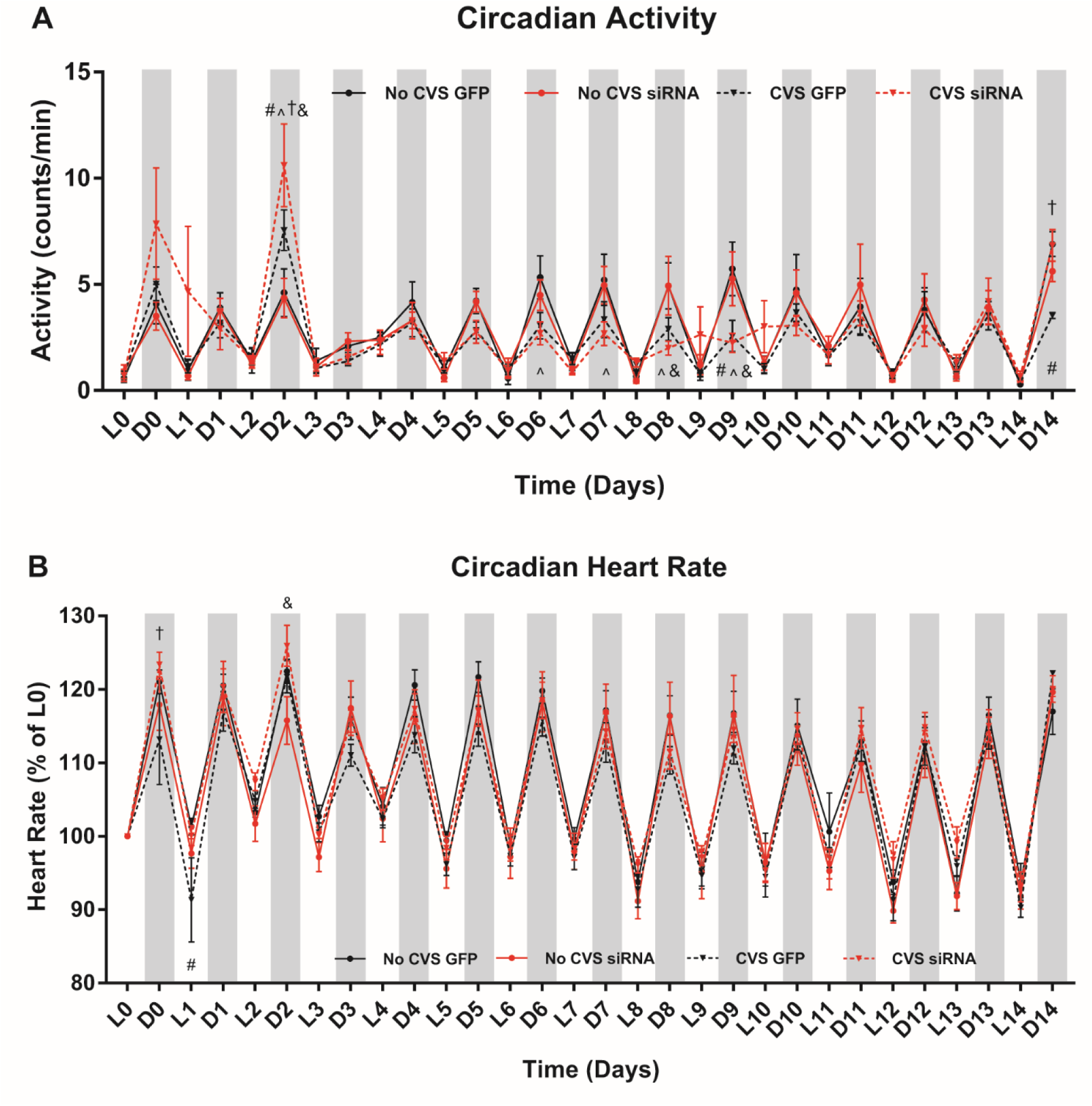
Chronically stressed rats had disrupted circadian behavioral rhythms evidenced by interactions of CVS and siRNA, as well as CVS and time (n = 7-8/group) (A). Circadian heart rate also exhibited a CVS x siRNA interaction (n = 8/group) leading to heart rate disruptions early in CVS (B). ^#^p < 0.05 CVS GFP vs. No CVS GFP, ^p < 0.05 CVS siRNA vs. No CVS GFP, ^†^p < 0.05 CVS siRNA vs. CVS GFP, ^&^p < 0.05 CVS siRNA vs. No CVS siRNA.

##### Circadian heart rate

Home cage HR (n = 8/group) was analyzed by 3-way repeated-measures ANOVA revealing a main effect of time, [F(29, 960) = 72.64, p < 0.0001] accompanied by a stress x treatment interaction [F(1, 960) = 28.71, p < 0.0001]. Early in CVS (Fig. 2B), the CVS siRNA group had elevated dark phase HR compared to the CVS GFP group (Day 0, p = 0.024), as well as the No siRNA group during overnight social crowding (Day 2, p = 0.024). Also, on the first light period of CVS, the CVS GFP HR was decreased compared to No CVS GFP (p = 0.028).

##### Circadian and cumulative arterial pressures

In order to examine the effects of chronic stress and decreased IL output on long-term blood pressure regulation, home cage arterial pressures (n = 6-8/group) were continuously monitored. For mean arterial pressure (MAP), 3-way repeated-measures ANOVA found main effects of stress [F(1, 900) = 15.33, p < 0.0001] and time [F(29, 900) = 13.91, p < 0.0001], as well as an interaction of stress x treatment [F(1, 900) = 14.41, p < 0.0001]. In terms of time-specific effects, there was an increase in MAP in the CVS siRNA group compared to No CVS siRNA (p = 0.022, Fig. 3A) during the light period of CVS day 1. While few time point-specific circadian effects were identified, AUC analysis found that the CVS siRNA group experienced greater cumulative MAP (p < 0.0001, Fig. 3B) and systolic arterial pressure (SAP, p < 0.0001, Fig. 3B) than all other groups. Diastolic arterial pressure (DAP, p < 0.0001, Fig. 3D) was also elevated in the CVS siRNA animals compared to No CVS siRNA and CVS GFP. Between non-stressed rats, the siRNA treatment led to lower cumulative pressures (MAP, SAP, and DAP, p < 0.0001) relative to GFP.

**Figure 3.**
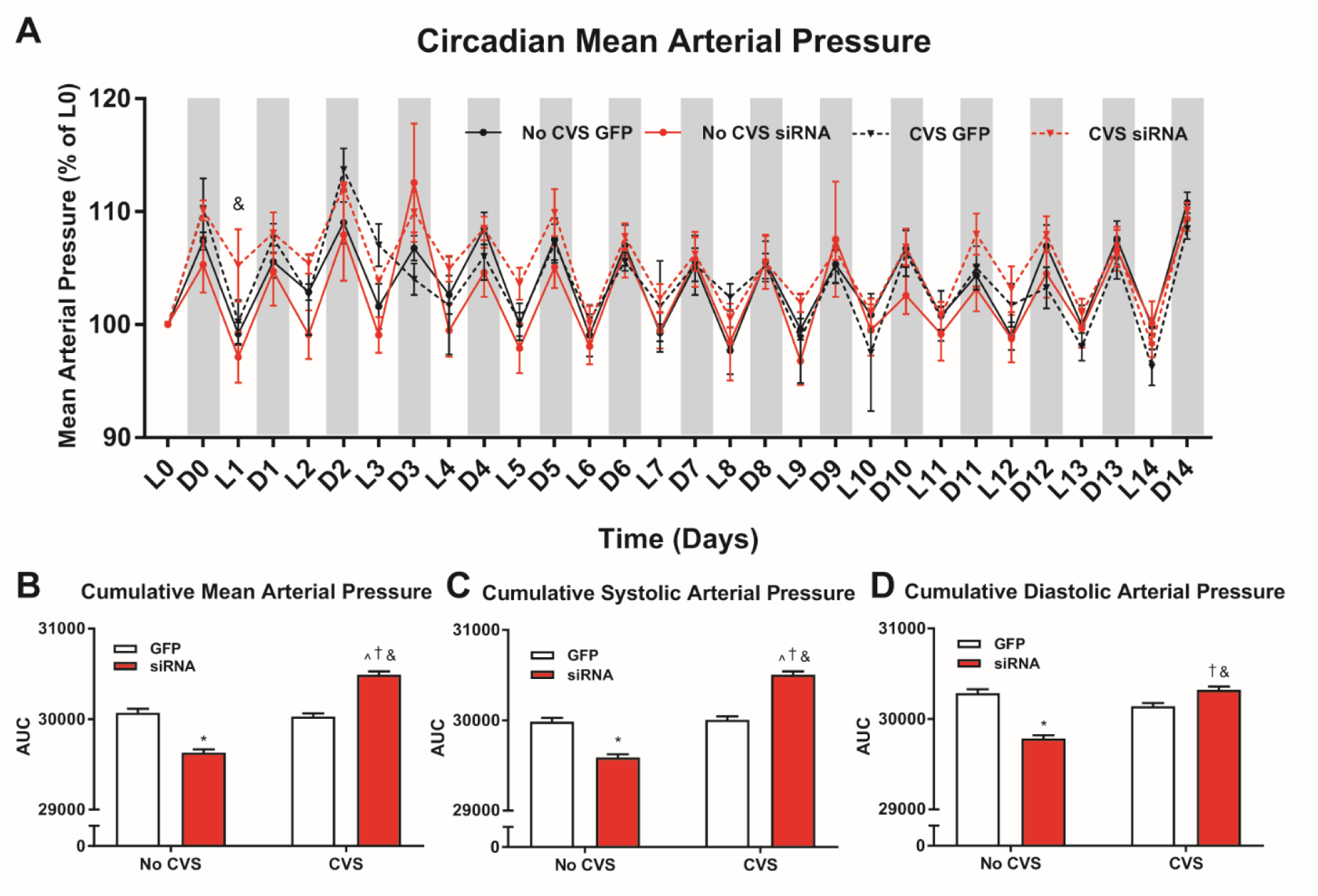
Chronic stress and siRNA treatment (n = 8/group) interacted leading to altered circadian arterial pressure (A). Analysis of cumulative arterial pressure indicated that CVS, only in siRNA-treated rats, increased chronic MAP (b), SAP (c), and DAP (d). *p < 0.05 No CVS siRNA vs. No CVS GFP, ^p < 0.05 CVS siRNA vs. No CVS GFP, ^†^p < 0.05 CVS siRNA vs. CVS GFP, ^&^p < 0.05 CVS siRNA vs. No CVS siRNA.

##### Circadian and cumulative pulse pressure

Pulse pressure is a function of vascular stiffness and predicts heart disease independent of MAP^41, 42^. Three-way repeated-measures ANOVA identified a main effect of stress [F (1, 900) = 34.35, p < 0.0001] and an interaction of stress x treatment [F(29, 900 = 2.59, p < 0.0001] for circadian pulse pressure. Over the course of CVS, the only time-specific difference in circadian pulse pressure occurred during the dark phase of day 0 (Fig. 4A) where the CVS GFP pulse pressure was greater than No CVS GFP (p = 0.01, n = 6-8/group). Cumulative pulse pressure from AUC analysis was increased in both CVS groups relative to No CVS (p < 0.0001, Fig 4b). Additionally, CVS siRNA cumulative pulse pressure was greater than all groups, including CVS GFP (p < 0.0001).

**Figure 4.**
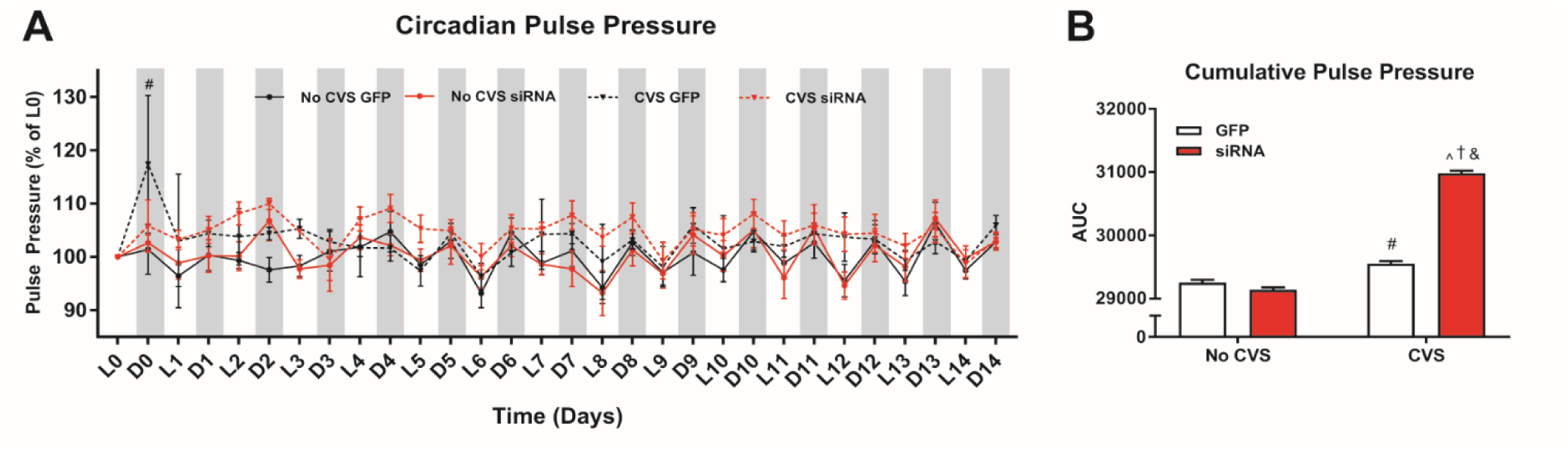
Chronic stress and siRNA treatment (n = 6-8/group) interacted to affect circadian pulse pressure (A). Cumulative pulse pressure was increased in CVS exposed rats with CVS siRNA rats experiencing the greatest chronic pulse pressure (B). ^#^p < 0.05 CVS GFP vs. No CVS GFP, ^p < 0.05 CVS siRNA vs. No CVS GFP, ^†^p < 0.05 CVS siRNA vs. CVS GFP, ^&^p < 0.05 CVS siRNA vs. No CVS siRNA.

##### Acute stress reactivity

In order to study the role of the IL in acute stress reactivity, MAP and HR reactivity were monitored during restraint (n = 8/group). During acute stress, 3-way repeated-measures ANOVA of HR reactivity found a main effect of time [F(1, 84) = 53.99, p < 0.0001] (Fig. 5A). Compared to No CVS GFP, vGluT1 siRNA elevated HR during restraint minutes 15-25 (p ≤ 0.040). Both CVS GFP and CVS siRNA rats had elevated HR compared to No CVS GFP from 10-25 minutes of restraint (p ≤ 0.006). Further, CVS siRNA rats had elevated HR during restraint at 35 and 40 minutes (p ≤ 0.008). While recovering from stress in the home cage, HR remained elevated in the CVS siRNA group relative to No CVS GFP at minutes 45 and 95 (p ≤ 0.010).

**Figure 5.**
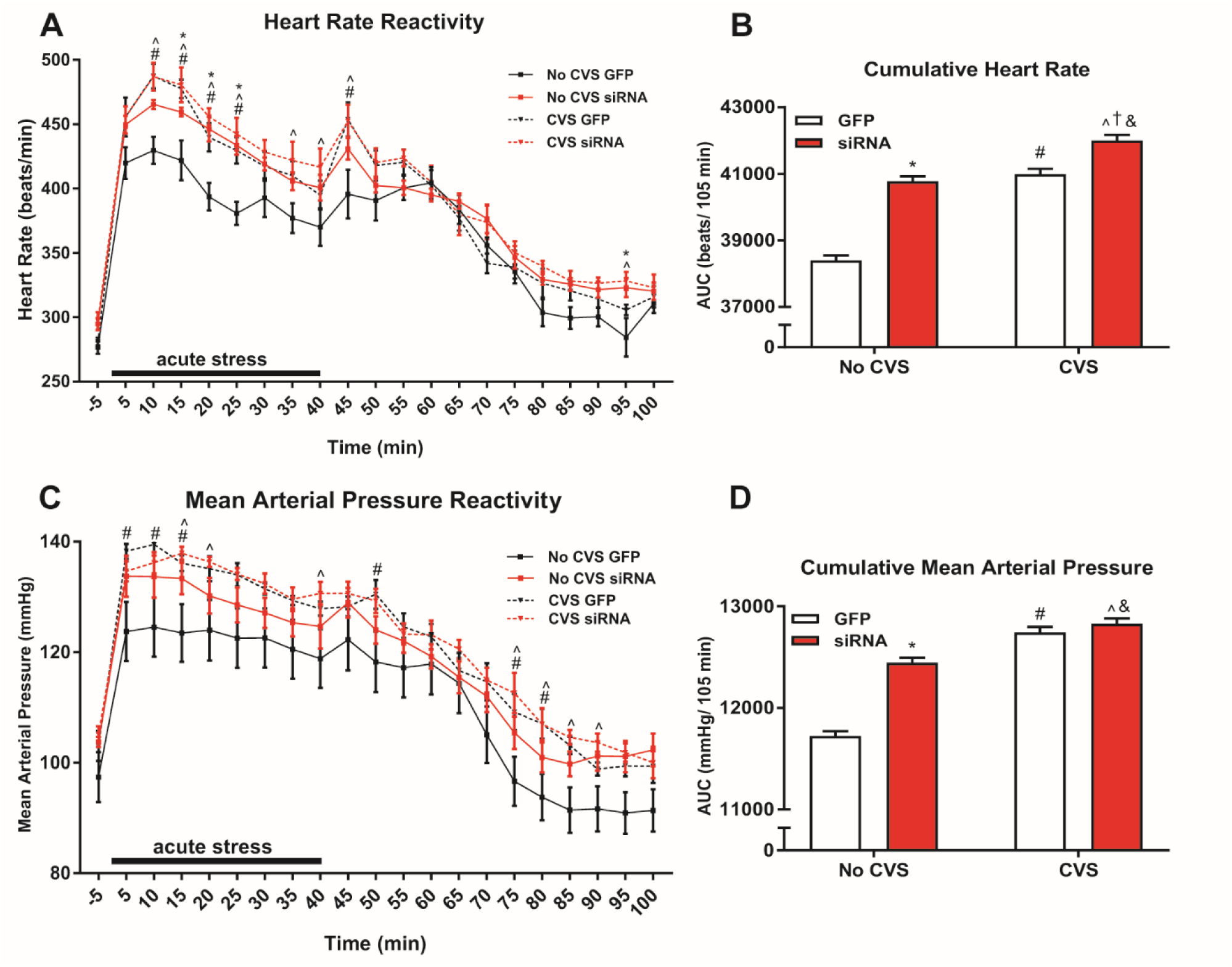
In response to acute restraint, both siRNA and CVS (n = 8/group) increased heart rate reactivity and impaired recovery (A). Cumulative acute heart rate responses were also elevated by siRNA and CVS but CVS siRNA rats had the greatest overall heart rate response (B). Chronically stressed rats, both GFP and siRNA treated, had increased MAP reactivity and impaired recovery (C). Analysis of cumulative restraint-induced pressor responses indicated effects of both siRNA and CVS (D). *p < 0.05 No CVS siRNA vs. No CVS GFP, ^#^p < 0.05 CVS GFP vs. No CVS GFP, ^p < 0.05 CVS siRNA vs. No CVS GFP, ^†^p < 0.05 CVS siRNA vs. CVS GFP, ^&^p < 0.05 CVS siRNA vs. No CVS siRNA.

Additionally, CVS GFP and No CVS siRNA had elevated HR on minutes 45 (p = 0.0003) and 95 (p = 0.031), respectively. Cumulative HR reactivity from AUC analysis of acute stress responses revealed that both No CVS siRNA and CVS GFP groups had elevated HR responses to acute restraint (p < 0.0001, Fig. 5B). Moreover, the CVS siRNA group experienced greater cumulative HR than all other groups (p = 0.0006).

Analysis of stress-evoked MAP by 3-way repeated-measures ANOVA identified a main effect of time [F(1, 84) = 99.91, p < 0.0001] (Fig. 5c). The CVS GFP group had greater MAP reactivity compared to No CVS GFP on minutes 5-15 of restraint (p ≤ 0.030). The CVS siRNA animals had greater MAP than No CVS GFP at 15, 20, and 40 minutes (p ≤ 0.05). During recovery, CVS GFP MAP remained elevated at 50, 75, and 80 minutes (p ≤ 0.04); furthermore, CVS siRNA MAP was higher at minutes 75-90 (p ≤ 0.045). AUC analysis by 2-way ANOVA found siRNA treatment increased cumulative MAP (p < 0.0001, Fig. 5D). CVS also increased MAP AUC as both CVS groups were higher than respective No CVS controls (p < 0.0001).

#### Experiment 2

##### Injection placement

Similar to experiment 1, injections of lentiviral-packaged constructs were targeted to the IL with minimal spread to PL (Fig. 6). Although, injections from experiment 2 had greater spread into superficial layers of the IL. Additionally, some injections spread into the striatum caudally but this region does not exhibit vGluT1 expression^43^.

**Figure 6.**
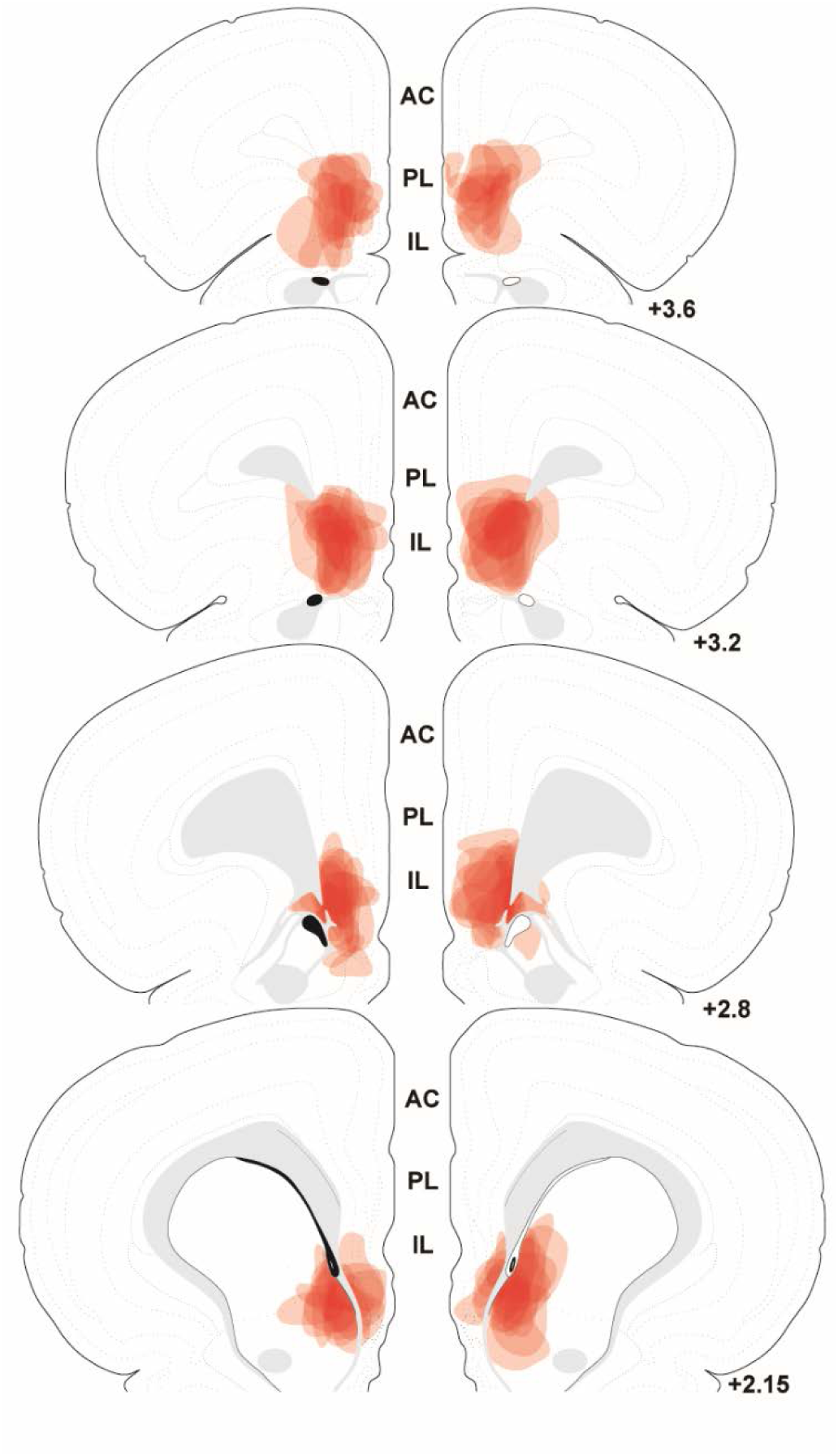
The spread of individual lentiviral injections was traced on photomicrographs and overlaid onto atlas templates from Swanson^74^ to depict the localization of vGluT1 knockdown in experiment 2. White arrows indicate dorsal and ventral boundaries of the IL. Numbers indicate distance rostral to bregma in millimeters. AC: anterior cingulate, PL: prelimbic cortex, IL: infralimbic cortex.

##### Vascular function

In order to assess the vascular consequences of prolonged stress, arterial function was monitored in response to endothelial-dependent and -independent agents *ex vivo*. In terms of endothelial-dependent vasoconstriction, 3-way repeated-measures ANOVA identified a main effect of KCl concentration [F(1,36) = 669.73, p < 0.0001] and an interaction of drug concentration x viral treatment [F(1,36) = 9.55, p = 0.004] (Fig. 7A). As determined by post-hoc analyses, aortas of CVS siRNA animals constricted less than No CVS GFP at KCl concentrations above 25 mM (p ≤ 0.0006). CVS siRNA also showed impaired constriction compared to CVS GFP at concentrations above 30 mM (p ≤ 0.006). Compared to No CVS siRNA, CVS siRNA vasoreactivity was decreased at concentrations of 30 and 50 mM (p ≤ 0.042). Within the No CVS groups, siRNA treatment decreased constriction at 40 mM (p = 0.019). Endothelium-independent vasoconstriction in response to phenylephrine showed a main effect of concentration [F(1,40) = 31.62, p < 0.0001] (Fig. 7B). Either siRNA or CVS alone impaired vasoreactivity at drug concentrations above 1 μM (p < 0.05). However, CVS siRNA animals had impaired vasoconstriction compared to all other groups at phenylephrine concentrations above 0.1 μM (p < 0.05).

**Figure 7.**
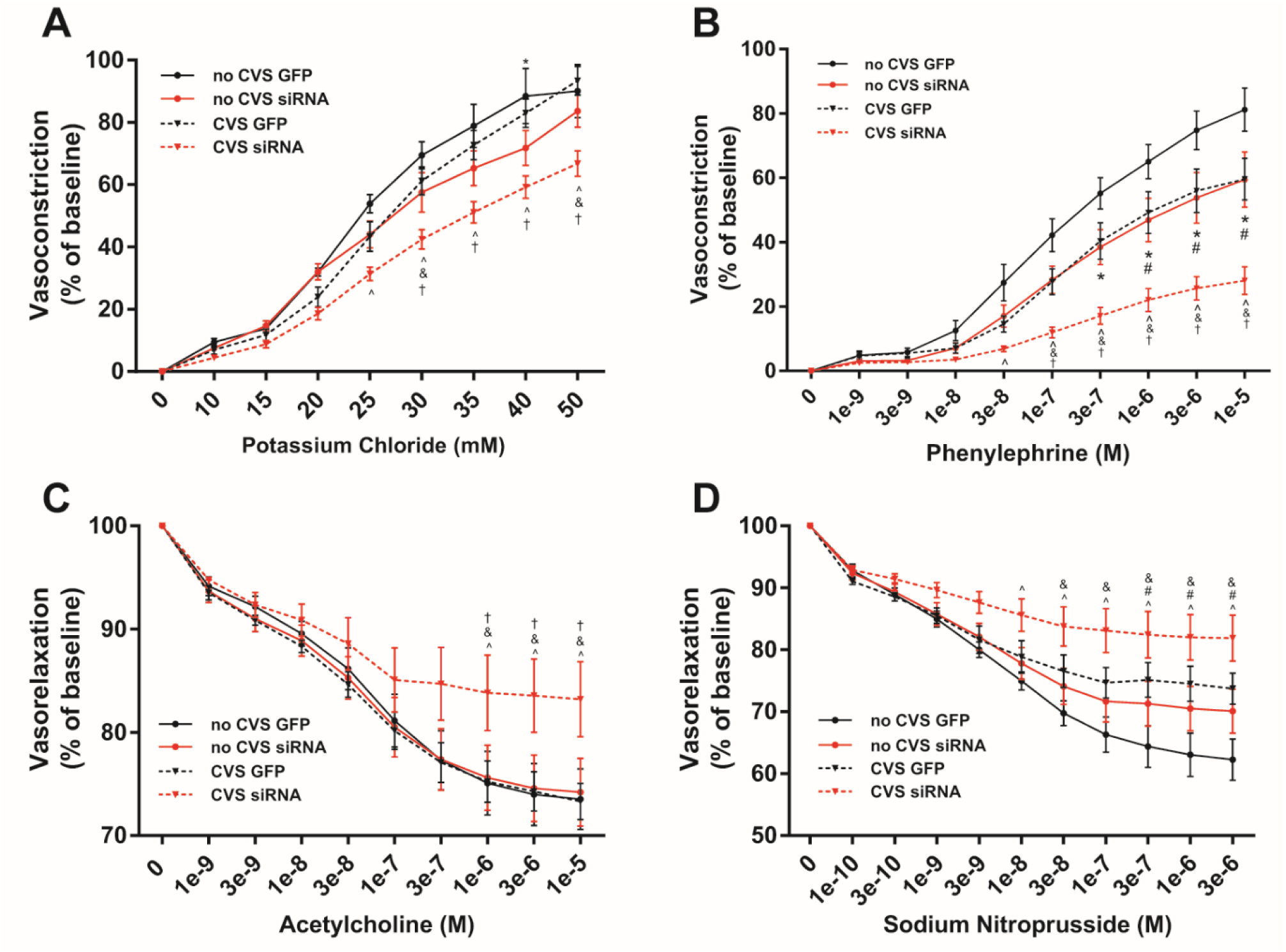
Aortic tissue from CVS siRNA animals (n = 7/group) had impaired endothelial-dependent vasoconstriction (A). Both CVS and siRNA impaired endothelial-independent vasoreactivity but the CVS siRNA group had the greatest impairment (B). Endothelial-dependent vasorelaxation was impaired only in the CVS siRNA group (C), while endothelial-independent relaxation was impaired in CVS GFP and CVS siRNA tissue (D). *p < 0.05 No CVS siRNA vs. No CVS GFP, ^#^p < 0.05 CVS GFP vs. No CVS GFP, ^p < 0.05 CVS siRNA vs. No CVS GFP, ^†^p < 0.05 CVS siRNA vs. CVS GFP, ^&^p < 0.05 CVS siRNA vs. No CVS siRNA.

Endothelial-dependent vasorelaxation to acetylcholine showed a main effect of drug concentration [F(1,40) = 15.89, p = 0.0003] by 3-way repeated-measures ANOVA (Fig. 7C). Post-test found that CVS siRNA vasorelaxation was decreased compared to all groups at acetylcholine concentrations above 1 μM (p = 0.048). Endothelial-independent vasorelaxation to SNP, as analyzed by 3-way repeated-measures ANOVA, showed a main effect of SNP concentration [F(1,44) = 12.65, p = 0.0009] (Fig. 7D). CVS siRNA tissue had impaired vasorelaxation compared to both No CVS groups at concentrations above of 0.03 μM (p = 0.026). Additionally, CVS alone impaired vasorelaxation in GFP rats at SNP concentrations above 0.3 μM (p < 0.010).

##### Vascular and cardiac histology

Histological analysis was carried out to investigate the effects of chronic stress and vGluT1 knockdown on markers of vascular and cardiac pathology (Table 2). In GFP-injected rats, CVS increased aortic tunica media thickness [F(1,24) = 18.55, p = 0.0002] and media:lumen area [F(1,24) = 10.45, p < 0.004]. Furthermore, CVS increased adventitial fibrosis in terms of increased collagen density [F(1,24) = 4.944, p = 0.036]. Chronic stress also affected the myocardium by increasing heart weight [F(1,24) = 7.028, p = 0.014] and myocyte surface area [F(1,23) = 5.084, p = 0.034]. In animals with reduced vGluT1, CVS had greater effects on aortic remodeling. CVS siRNA rats had decreased luminal circumference [F(1,24) = 8.217, p = 0.022] and area [F(1,24) = 8.142, p = 0.026], increased media thickness [F(1,24) = 18.55, p = 0.026] and media:lumen area [F(1,24) = 10.45, p < 0.0004], increased collagen density [F(1,24) = 4.944, p = 0.036] and decreased adventitial thickness [F(1,24) = 6.517, p = 0.031] (Fig. 8A-D). Collectively, these results indicate that CVS interacts with decreased IL function to promote fibrosis and inward remodeling of vascular muscle leading to restricted luminal area, potentially accounting for impaired vasoreactivity and arterial stiffness. CVS siRNA rats also exhibited cardiac hypertrophy as these animals had increased heart weight [F(1,24) = 7.028, p = 0.014], heart weight relative to body weight [F(1,24) = 17.09, p = 0.015], and increased myocyte surface area [F(1,23) = 5.084, p = 0.034] (Fig. 8E, F), without affecting myocardial collagen deposition.

**Figure 8.**
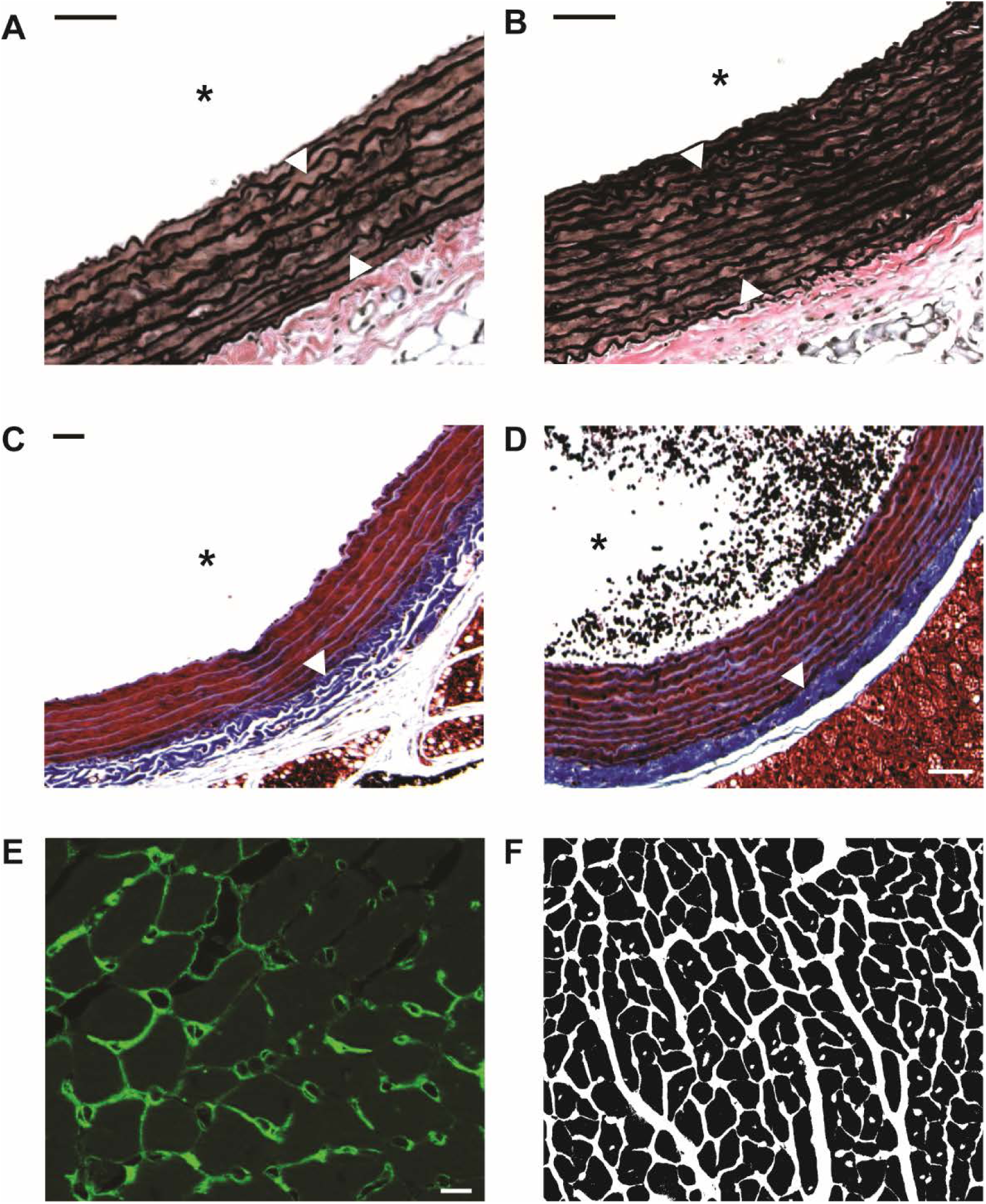
Verhoeff-van Gieson stain was used to visualize elastin (dark brown) in aortic tissue of No CVS GFP (A) and CVS siRNA (B) rats. Greater thickness of the tunica media is indicated by white arrows. Masson’s Trichrome was used to stain collagen (blue) in aortic tissue of No CVS GFP (C) and CVS siRNA (D) animals. White arrows indicate increased collagen density in the tunica adventitia. Wheat germ agglutinin conjugated to Alexa 488 (green) was used to visualize cardiomyocyte membranes (E). Binarized myocyte images (F) were used to quantify myocyte surface area. Scale bars: (A-D) 50 µm, (E) 10 µm. **\*** denotes the lumen.

**Table 2.** Vascular and cardiac histological analysis. Histological quantification of vascular and myocardial structure. Dimensions of the lumen and media were quantified from elastin staining. Adventitial size and fibrosis were determined from collagen staining. Myocyte surface area was measured with membrane labeling and cardiac fibrosis was queried via collagen staining. There were no effects of siRNA alone. CVS increased media thickness, adventitial collagen, and myocyte size. In contrast, CVS siRNA tissue exhibited alterations in all structural endpoints assessed except myocardial fibrosis.

| **Table 2. Vascular and cardiac histological analysis** |  |  |  |  |
| --- | --- | --- | --- | --- |
| **n = 7/group** | **No CVS GFP** | **No CVS siRNA** | **CVS GFP** | **CVS siRNA** |
| **Luminal Circumference (mm)** | 4.967 ± 0.058 | 5.135 ± 0.058 | 4.897 ± 0.069 | 4.893 ± 0.019<sup>&</sup> |
| **Luminal Area (mm<sup>2</sup>)** | 1.694 ± 0.035 | 1.786 ± 0.037 | 1.619 ± 0.037 | 1.550 ± 0.089<sup>&</sup> |
| **Media Thickness (mm)** | 0.107 ± 0.001 | 0.108 ± 0.002 | 0.113 ± 0.001<sup>#</sup> | 0.114 ± 0.001<sup>^&</sup> |
| **Media:Lumen Area** | 0.341 ± 0.009 | 0.335 ± 0.014 | 0.381 ± 0.005<sup>#</sup> | 0.392 ± 0.024<sup>^&</sup> |
| **Adventitia Collagen (% area)** | 66.93 ± 1.320 | 66.82 ± 0.893 | 71.49 ± 2.313<sup>#</sup> | 70.86 ± 2.659<sup>^&</sup> |
| **Adventitia Thickness** | 0.040 ± 0.001 | 0.037 ± 0.002 | 0.036 ± 0.002 | 0.034 ± 0.001<sup>^</sup> |
| **Heart Weight (g)** | 1.264 ± 0.018 | 1.266 ± 0.038 | 1.321 ± 0.028<sup>#</sup> | 1.365 ± 0.029<sup>^&</sup> |
| **Heart Weight/Body Weight (x100)** | 0.335 ± 0.005 | 0.340 ± 0.009 | 0.355 ± 0.007 | 0.366 ± 0.007<sup>^</sup> |
| **Myocyte Surface Area (mm<sup>2</sup>)** | 349.9 ± 18.70 | 352.1 ± 11.66 | 396.9 ± 18.84<sup>#</sup> | 406.8 ± 28.53<sup>^&</sup> |
| **Myocardial Collagen (% area)** | 0.429 ± 0.013 | 0.449 ± 0.023 | 0.479 ± 0.042 | 0.459 ± 0.016 |

<sup>#</sup>p < 0.05 CVS GFP vs. No CVS GFP, <sup>^</sup>p < 0.05 CVS siRNA vs. No CVS GFP, <sup>&</sup>p < 0.05 CVS siRNA vs. No CVS siRNA

## Discussion

In the current study, we utilized viral-mediated gene transfer to decrease IL glutamatergic output while simultaneously monitoring hemodynamic, vascular, and cardiac responses to chronic stress. We found that IL vGluT1 knockdown increased heart rate and arterial pressure reactivity to acute stress. Additionally, IL hypofunction during CVS increased chronic home cage arterial pressure. These changes were accompanied by both endothelial-independent and -dependent arterial dysfunction. Histological analysis revealed that animals experiencing CVS with decreased IL output also had inward vascular remodeling, fibrosis, and cardiac hypertrophy. Collectively, these results indicate that IL projection neurons are critical for reducing acute cardiovascular stress reactivity, long-term arterial pressure, and vascular endothelial dysfunction. Furthermore, they identify a neurochemical mechanism linking stress appraisal and emotion with chronic stress-induced autonomic dysfunction.

Epidemiological evidence indicates that prolonged stress is a major risk factor for cardiovascular illness and mortality^7, 44^. Additionally, numerous clinical studies point to enhanced stress reactivity as a marker of future cardiovascular pathology^9, 45^. Rodent studies employing repeated-stress models of depression (chronic variable stress, chronic mild stress, chronic social defeat, etc.) have found alterations in baroreflex function, decreased heart rate variability, and ventricular arrhythmias^4, 46–52^. Given the strong association between prolonged stress/emotional disorders and cardiovascular disease^53–55^, it is important to identify the specific neural processes of stress appraisal and mood that impact cardiovascular function. Emerging evidence suggests that autonomic imbalance prolongs exposure to neural and endocrine stress mediators, generating risk for cardiovascular pathology^56–58^. Stress-associated molecules such as corticosteroids, corticotropin-releasing hormone, and neuropeptide Y, among others, have been shown to affect cardiovascular function in animal models^50, 59, 60^. However, the neural circuits that integrate cognitive appraisal and autonomic balance remain to be determined.

The contribution of the current study relates to the site-specific genetic approach that reduces IL vGluT1 expression long-term^28^. Given the decreased output from a critical cognitive/emotional region^11, 32^, we monitored both stress-induced and resting parameters of heart rate and arterial pressure in otherwise unstressed rats, as well as rats experiencing the cumulative burden of chronic stress exposure. This approach was followed by *ex vivo* analysis of arterial function and histological investigation of vascular and myocardial structure. These studies found several siRNA effects in animals that did not experience CVS (Table 3). The knockdown increased HR and MAP reactivity to acute restraint and impaired endothelial-independent vasoconstriction, suggesting hypo-function of IL primes for enhanced cardiovascular reactivity and impaired vasomotor function. Previous studies indicating that NMDA-mediated activation of the IL reduces tachycardic and pressor responses to acute air-jet stress further support the conclusion that IL glutamate output reduces acute cardiovascular reactivity^61^. Paradoxically, No CVS siRNA animals exhibited decreased resting home cage circadian arterial pressure. This effect may relate to the important role of IL molecular clocks in coordinating circadian physiological rhythms^62^. In GFP-treated rats, CVS dampened home cage activity in the dark cycle, decreasing circadian rhythms of activity and potentially accounting for effects of CVS to decrease circadian HR. Interestingly, CVS did not alter chronic home cage MAP; however, CVS-exposed animals exhibited enhanced tachycardic and pressor responses to acute restraint. This was accompanied by impaired endothelial-independent vasorelaxation and constriction. Furthermore, CVS increased vascular smooth muscle thickness and fibrosis, as well as cardiomyocyte surface area.

**Table 3.**
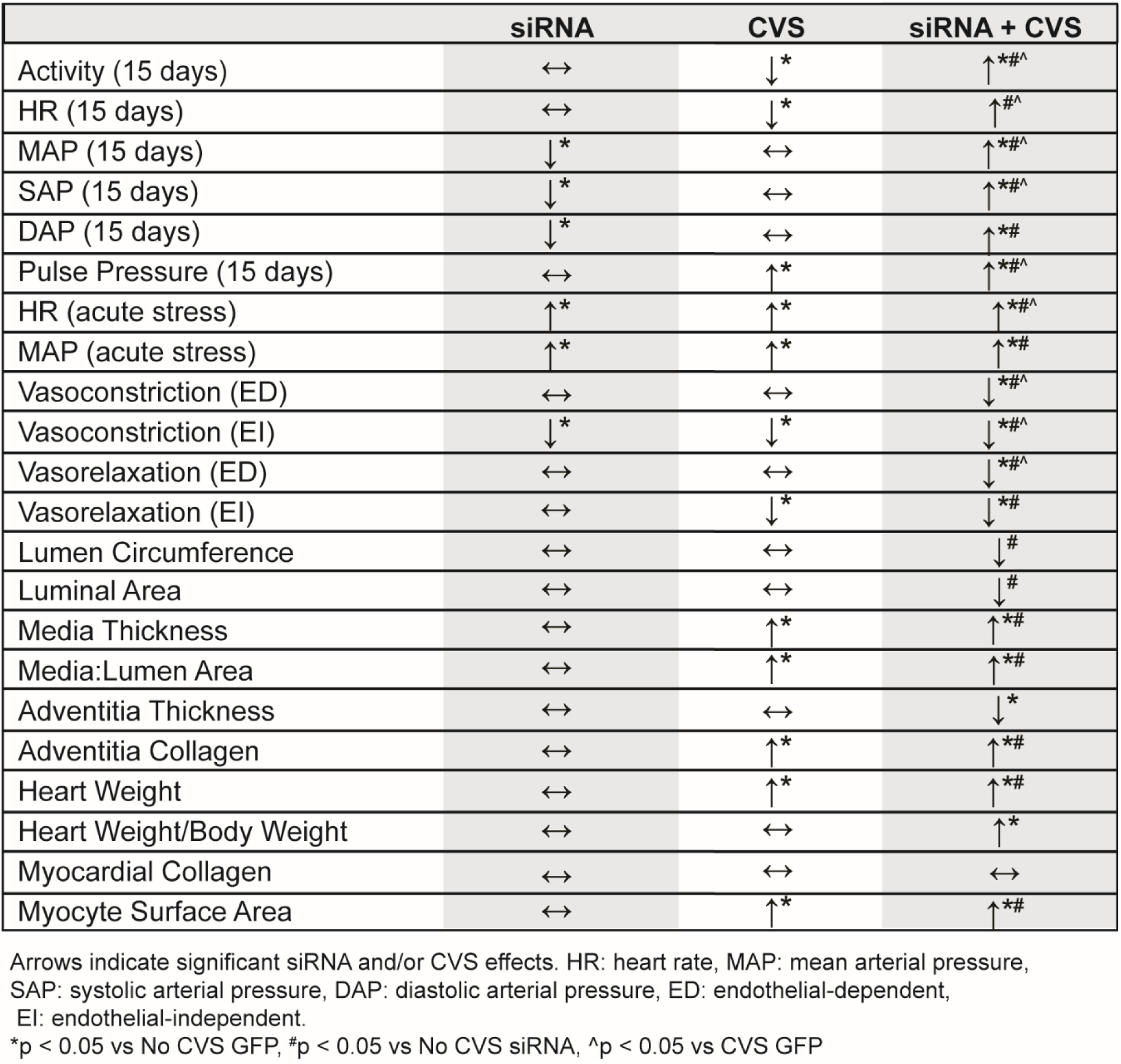
Summary: siRNA and/or CVS effects from experiments 1 and 2. Integrative summary of all data reported in terms of siRNA effects, chronic stress effects, and the combination of siRNA and CVS.

The effects of vGluT1 knockdown and CVS interacted to generate a phenotype of enhanced cardiovascular risk. All measures of HR and arterial pressure, both acute stress-induced and chronic home cage, were elevated. This group also had impairment of both endothelial-dependent and –independent vascular dilation and constriction. The overall risk profile was further evident by inward remodeling of the vasculature due to hypertrophy and fibrosis that reduced luminal area, indicative of vascular stiffness. Additionally, these rats exhibited myocardial hypertrophy including increased myocyte size, heart weight, and body weight-corrected heart weight. It is worth noting that some effects in the CVS siRNA group result from comparisons to No CVS GFP. In the case of histological measures, there were no significant differences between CVS GFP and CVS siRNA animals. This suggests that the structural changes observed are not sufficient to fully account for vascular dysfunction in the CVS siRNA group relative to CVS GFP. Furthermore, home cage arterial pressure elevations were modest and not indicative of a hypertensive state. More likely, endothelial dysfunction after chronic stress in animals with impaired IL function results from the interaction of multiple factors, including enhanced HPA axis activity, elevated cardiovascular reactivity, increased resting arterial pressure, and vascular remodeling. Taken together, these results suggest that conditions associated with decreased ventral mPFC activity, including depression, anxiety, and post-traumatic stress disorder^32^, may enhance vulnerability to the effects of prolonged stress on cardiovascular health.

The specific circuit mechanisms downstream of IL glutamate release that account for the current findings remain to be determined. Although the IL does not directly innervate either pre-ganglionic autonomic neurons or neurosecretory cells of the hypothalamus^21, 63^, the region has widespread projections throughout the forebrain and brainstem^21, 25^. Inputs to the posterior hypothalamus target local inhibitory cells^23^, providing a potential pathway to inhibit stress reactivity^64^. Additionally, IL-targeted neurons of the PH innervate the paraventricular hypothalamus, a region important for vasomotor tone^23, 65^. There are also IL projections to brainstem cardioregulatory centers including the nucleus of the solitary tract and the rostral and caudal regions of the ventrolateral medulla that may regulate autonomic outflow^24, 66^. Furthermore, there is a complex network of local cortical circuitry, including interneurons in the IL, that mediates the overall activity of glutamatergic projection neurons^67^. Based on the current findings, these local circuits would be expected to play a role in autonomic reactivity. Ultimately, projection-specific analyses are needed isolate the precise cell populations within IL glutamate neurons that reduce autonomic imbalance after chronic stress.

While these studies identified a novel frontal cortical node for preventing the deleterious cardiovascular effects of chronic stress, there are limitations worth discussing. First, the current studies were limited to males. As depression-cardiovascular co-morbidity has at least twice the prevalence in females^68–70^, it is important to consider sex-specific regulation. Interestingly, a recent study investigating vascular function in female rodents after chronic stress actually found that ovarian hormones protect against stress-induced arterial dysfunction^71^. Comparing the neural basis of pathological responses in males and females would likely yield a better understanding of the disproportionate female impact of mood disorder-cardiovascular co-morbidity. Another consideration is that the current vasoreactivity experiments were carried out in thoracic aorta. Although the results identified impaired function and indicators of stiffness, the aorta is a conductive artery and future experiments with resistance arterioles might yield differing results. Indeed, these studies could show greater effects as resistance arteries receive more sympathetic innervation^72, 73^. Furthermore, investigating vascular resistance could yield data with significant relevance for stress-related hypertension.

In conclusion, the current findings highlight IL glutamatergic neurons as a node of integration that links stress appraisal with hemodynamic reactivity, long-term arterial pressure control, and vascular endothelial function. These results also indicate that, in the context of chronic stress, cortical cells mediating cognition and behavior can impact the structure and function of the heart and vasculature. Future research investigating the mechanisms that regulate IL projection neuron activity and the downstream post-synaptic events activated by IL glutamate release may yield insight into novel targets to prevent or reduce the burden of cardiovascular disease.

## Acknowledgments

This work was supported by NIH grants K99/R00 HL122454 to B. Myers and R01 MH049698 to J. P. Herman. Cincinnati Children’s Hospital Medical Center Research Pathology Core carried out cardiac and vascular histological processing.

## Disclosures

J. M. McKlveen contributed to this article in her personal capacity. The views expressed herein are those of the authors and do not necessarily represent the views of the National Institutes of Health, National Center for Complementary and Integrative Health, or the United States Government. All authors report no biomedical financial interests or potential conflicts of interest.

This article was first published as a preprint: Schaeuble D, Packard AEB, McKlveen JM, Morano R, Fourman S, Smith BL, Scheimann JR, Packard B, Wilson SP, James J, Hui D, Ulrich-Lai YU, Herman JP, Myers B. Prefrontal cortical regulation of chronic stress-induced cardiovascular susceptibility. *bioRxiv*. doi.org/10.1101/675835.

## References

1. de Kloet ER, Joëls M, Holsboer F. Stress and the brain: from adaptation to disease. Nat Rev Neurosci. 2005;6:463–475.

2. Myers B, McKlveen JM, Herman JP. Glucocorticoid actions on synapses, circuits, and behavior: Implications for the energetics of stress. Front Neuroendocrinol. 2014;35:180–196.

3. Wardle J, Chida Y, Gibson EL, Whitaker KL, Steptoe A. Stress and Adiposity: A Meta-Analysis of Longitudinal Studies. Obesity. 2011;19:771–778.

4. Grippo AJ, Johnson AK. Stress, depression and cardiovascular dysregulation: a review of neurobiological mechanisms and the integration of research from preclinical disease models. Stress. 2009;12:1–21.

5. Binder EB, Nemeroff CB. The CRF system, stress, depression and anxiety— insights from human genetic studies. Mol Psychiatry. 2010;15:574–588.

6. Sgoifo A, Carnevali L, Pico Alfonso MDLA, Amore M. Autonomic dysfunction and heart rate variability in depression. Stress. 2015;18:343–352.

7. Yusuf S, Hawken S, Ôunpuu S, Dans T, Avezum A, Lanas F, McQueen M, Budaj A, Pais P, Varigos J, Lisheng L. Effect of potentially modifiable risk factors associated with myocardial infarction in 52 countries (the INTERHEART study): case-control study. Lancet. 2004;364:937–952.

8. Barefoot JC, Helms MJ, Mark DB, Blumenthal JA, Califf RM, Haney TL, O’Connor CM, Siegler IC, Williams RB. Depression and Long-Term Mortality Risk in Patients With Coronary Artery Disease. Am J Cardiol. 1996;78:613–617.

9. Chida Y, Steptoe A. Greater Cardiovascular Responses to Laboratory Mental Stress Are Associated With Poor Subsequent Cardiovascular Risk Status. Hypertension. 2010;55:1026–1032.

10. Myers-Schulz B, Koenigs M. Functional anatomy of ventromedial prefrontal cortex: implications for mood and anxiety disorders. Mol Psychiatry. 2012;17:132– 141.

11. McKlveen JM, Myers B, Herman JP. The Medial Prefrontal Cortex: Coordinator of Autonomic, Neuroendocrine and Behavioural Responses to Stress. J Neuroendocrinol. 2015.

12. Damasio AR. The somatic marker hypothesis and the possible functions of the prefrontal cortex. Philos Trans R Soc Lond B Biol Sci. 1996;351:1413–20.

13. Wood JN, Grafman J. Human prefrontal cortex: processing and representational perspectives. Nat Rev Neurosci. 2003;4:139–147.

14. Liotti M, Mayberg HS, Brannan SK, McGinnis S, Jerabek P, Fox PT. Differential limbic–cortical correlates of sadness and anxiety in healthy subjects: implications for affective disorders. Biol Psychiatry. 2000;48:30–42.

15. Mayberg HS, Lozano AM, Voon V, McNeely HE, Seminowicz D, Hamani C, Schwalb JM, Kennedy SH. Deep Brain Stimulation for Treatment-Resistant Depression. Neuron. 2005;45:651–660.

16. Shoemaker JK, Badrov MB, Al-Khazraji BK, Jackson DN. Neural Control of Vascular Function in Skeletal Muscle. In: Comprehensive Physiology. Hoboken, NJ, USA: John Wiley & Sons, Inc.; 2015: 303–329.

17. Beissner F, Meissner K, Bär K-J, Napadow V. The Autonomic Brain: An Activation Likelihood Estimation Meta-Analysis for Central Processing of Autonomic Function. J Neurosci. 2013;33:10503–10511.

18. Gianaros PJ, Sheu LK. A review of neuroimaging studies of stressor-evoked blood pressure reactivity: emerging evidence for a brain-body pathway to coronary heart disease risk. Neuroimage. 2009;47:922–36.

19. Gianaros PJ, Wager TD. Brain-Body Pathways Linking Psychological Stress and Physical Health. Curr Dir Psychol Sci. 2015;24:313–321.

20. Uylings HBM, Groenewegen HJ, Kolb B. Do rats have a prefrontal cortex? Behav Brain Res. 2003;146:3–17.

21. Vertes RP. Differential projections of the infralimbic and prelimbic cortex in the rat. Synapse. 2004;51:32–58.

22. Öngür D, Ferry AT, Price JL. Architectonic subdivision of the human orbital and medial prefrontal cortex. J Comp Neurol. 2003;460:425–449.

23. Myers B, Carvalho-Netto E, Wick-Carlson D, Wu C, Naser S, Solomon MB, Ulrich-Lai YM, Herman JP. GABAergic Signaling within a Limbic-Hypothalamic Circuit Integrates Social and Anxiety-Like Behavior with Stress Reactivity. Neuropsychopharmacology. 2016;41:1530–1539.

24. Gabbott PLA, Warner TA, Jays PRL, Salway P, Busby SJ. Prefrontal cortex in the rat: Projections to subcortical autonomic, motor, and limbic centers. J Comp Neurol. 2005;492:145–177.

25. Wood M, Adil O, Wallace T, Fourman S, Wilson SP, Herman JP, Myers B. Infralimbic prefrontal cortex structural and functional connectivity with the limbic forebrain: a combined viral genetic and optogenetic analysis. Brain Struct Funct. 2019;224:73–97.

26. Myers B, Mark Dolgas C, Kasckow J, Cullinan WE, Herman JP. Central stress-integrative circuits: forebrain glutamatergic and GABAergic projections to the dorsomedial hypothalamus, medial preoptic area, and bed nucleus of the stria terminalis. Brain Struct Funct. 2014;219:1287–303.

27. Smith BL, Lyons CE, Correa FG, Benoit SC, Myers B, Solomon MB, Herman JP. Behavioral and physiological consequences of enrichment loss in rats. Psychoneuroendocrinology. 2017;77:37–46.

28. Myers B, McKlveen JM, Morano R, Ulrich-Lai YM, Solomon MB, Wilson SP, Herman JP. Vesicular Glutamate Transporter 1 Knockdown in Infralimbic Prefrontal Cortex Augments Neuroendocrine Responses to Chronic Stress in Male Rats. Endocrinology. 2017;158:3579–3591.

29. Schuske K, Jorgensen EM. Vesicular glutamate transporter--shooting blanks. Science. 2004;304:1750–2.

30. Wojcik SM, Rhee JS, Herzog E, Sigler A, Jahn R, Takamori S, Brose N, Rosenmund C. An essential role for vesicular glutamate transporter 1 (VGLUT1) in postnatal development and control of quantal size. Proc Natl Acad Sci U S A. 2004;101:7158–63.

31. Grillo CA, Piroli GG, Lawrence RC, Wrighten SA, Green AJ, Wilson SP, Sakai RR, Kelly SJ, Wilson MA, Mott DD, Reagan LP. Hippocampal Insulin Resistance Impairs Spatial Learning and Synaptic Plasticity. Diabetes. 2015;64:3927–36.

32. McKlveen JM, Myers B, Flak JN, Bundzikova J, Solomon MB, Seroogy KB, Herman JP. Role of Prefrontal Cortex Glucocorticoid Receptors in Stress and Emotion. Biol Psychiatry. 2013;74:672–679.

33. Grillo CA, Tamashiro KL, Piroli GG, Melhorn S, Gass JT, Newsom RJ, Reznikov LR, Smith A, Wilson SP, Sakai RR, Reagan LP. Lentivirus-mediated downregulation of hypothalamic insulin receptor expression. Physiol Behav. 2007;92:691–701.

34. Flak JN, Jankord R, Solomon MB, Krause EG, Herman JP. Opposing effects of chronic stress and weight restriction on cardiovascular, neuroendocrine and metabolic function. Physiol Behav. 2011;104:228–234.

35. Goodson ML, Packard AEB, Buesing DR, Maney M, Myers B, Fang Y, Basford JE, Hui DY, Ulrich-Lai YM, Herman JP, Ryan KK. Chronic stress and Rosiglitazone increase indices of vascular stiffness in male rats. Physiol Behav. 2017;172:16–23.

36. Flak JN, Solomon MB, Jankord R, Krause EG, Herman JP. Identification of chronic stress-activated regions reveals a potential recruited circuit in rat brain. Eur J Neurosci. 2012;36:2547–55.

37. Basford JE, Koch S, Anjak A, Singh VP, Krause EG, Robbins N, Weintraub NL, Hui DY, Rubinstein J. Smooth muscle LDL receptor-related protein-1 deletion induces aortic insufficiency and promotes vascular cardiomyopathy in mice. PLoS One. 2013;8:e82026.

38. Gupta MK, McLendon PM, Gulick J, James J, Khalili K, Robbins J. UBC9-Mediated Sumoylation Favorably Impacts Cardiac Function in Compromised Hearts. Circ Res. 2016;118:1894–905.

39. Bensley JG, De Matteo R, Harding R, Black MJ. Three-dimensional direct measurement of cardiomyocyte volume, nuclearity, and ploidy in thick histological sections. Sci Rep. 2016;6:23756.

40. Solomon MB, Jones K, Packard BA, Herman JP. The medial amygdala modulates body weight but not neuroendocrine responses to chronic stress. J Neuroendocrinol. 2010;22:13–23.

41. Glasser SP, Halberg DL, Sands C, Gamboa CM, Muntner P, Safford M. Is Pulse Pressure an Independent Risk Factor for Incident Acute Coronary Heart Disease Events? The REGARDS Study. Am J Hypertens. 2014;27:555–563.

42. Franklin SS, Khan SA, Wong ND, Larson MG, Levy D. Is pulse pressure useful in predicting risk for coronary heart Disease? The Framingham heart study. Circulation. 1999;100:354–60.

43. Ziegler DR, Cullinan WE, Herman JP. Distribution of vesicular glutamate transporter mRNA in rat hypothalamus. J Comp Neurol. 2002;448:217–229.

44. Steptoe A, Kivimäki M. Stress and cardiovascular disease. Nat Rev Cardiol. 2012;9:360–370.

45. Vogelzangs N, Beekman ATF, Milaneschi Y, Bandinelli S, Ferrucci L, Penninx BWJH. Urinary Cortisol and Six-Year Risk of All-Cause and Cardiovascular Mortality. J Clin Endocrinol Metab. 2010;95:4959–4964.

46. Grippo AJ, Moffitt JA, Johnson AK. Cardiovascular alterations and autonomic imbalance in an experimental model of depression. Am J Physiol Integr Comp Physiol. 2002;282:R1333–R1341.

47. Costoli T, Bartolomucci A, Graiani G, Stilli D, Laviola G, Sgoifo A. Effects of chronic psychosocial stress on cardiac autonomic responsiveness and myocardial structure in mice. Am J Physiol - Hear Circ Physiol. 2004;286:H2133–H2140.

48. Carnevali L, Trombini M, Rossi S, Graiani G, Manghi M, Koolhaas JM, Quaini F, Macchi E, Nalivaiko E, Sgoifo A. Structural and Electrical Myocardial Remodeling in a Rodent Model of Depression. Psychosom Med. 2013;75:42–51.

49. Wood SK. Cardiac autonomic imbalance by social stress in rodents: understanding putative biomarkers. Front Psychol. 2014;5:950.

50. Wood SK, McFadden K V., Grigoriadis D, Bhatnagar S, Valentino RJ. Depressive and cardiovascular disease comorbidity in a rat model of social stress: a putative role for corticotropin-releasing factor. Psychopharmacology (Berl). 2012;222:325– 336.

51. Duarte JO, Planeta CS, Crestani CC. Immediate and long-term effects of psychological stress during adolescence in cardiovascular function: Comparison of homotypic vs heterotypic stress regimens. Int J Dev Neurosci. 2015;40:52–59.

52. Crestani CC. Emotional Stress and Cardiovascular Complications in Animal Models: A Review of the Influence of Stress Type. Front Physiol. 2016;7:251.

53. Barton DA, Dawood T, Lambert EA, Esler MD, Haikerwal D, Brenchley C, Socratous F, Kaye DM, Schlaich MP, Hickie I, Lambert GW. Sympathetic activity in major depressive disorder: identifying those at increased cardiac risk? J Hypertens. 2007;25:2117–2124.

54. Carney RM, Blumenthal JA, Stein PK, Watkins L, Catellier D, Berkman LF, Czajkowski SM, O’Connor C, Stone PH, Freedland KE. Depression, Heart Rate Variability, and Acute Myocardial Infarction. Circulation. 2001;104:2024–2028.

55. Hausberg M, Hillebrand U, Kisters K. Addressing sympathetic overactivity in major depressive disorder. J Hypertens. 2007;25:2004–2005.

56. Johnson AK, Grippo AJ. Sadness and broken hearts: Neurohumoral mechanisms and co-morbidity of ischemic heart disease and psychological depression. In: Journal of Physiology and Pharmacology.; 2006: 5–29.

57. Thayer JF, Åhs F, Fredrikson M, Sollers JJ, Wager TD. A meta-analysis of heart rate variability and neuroimaging studies: Implications for heart rate variability as a marker of stress and health. Neurosci Biobehav Rev. 2012;36:747–756.

58. Wulsin LR, Horn PS, Perry JL, Massaro JM, D’Agostino RB. Autonomic Imbalance as a Predictor of Metabolic Risks, Cardiovascular Disease, Diabetes, and Mortality. J Clin Endocrinol Metab. 2015;100:2443–2448.

59. Oakley RH, Cruz-Topete D, He B, Foley JF, Myers PH, Xu X, Gomez-Sanchez CE, Chambon P, Willis MS, Cidlowski JA. Cardiomyocyte glucocorticoid and mineralocorticoid receptors directly and antagonistically regulate heart disease in mice. Sci Signal. 2019;12:eaau9685.

60. Costoli T, Sgoifo A, Stilli D, Flugge G, Adriani W, Laviola G, Fuchs E, Pedrazzini T, Musso E. Behavioural, neural and cardiovascular adaptations in mice lacking the NPY Y1 receptor. Neurosci Biobehav Rev. 2005;29:113–123.

61. Müller-Ribeiro FC de F, Zaretsky D V, Zaretskaia M V, Santos RAS, DiMicco JA, Fontes MAP. Contribution of infralimbic cortex in the cardiovascular response to acute stress. Am J Physiol Regul Integr Comp Physiol. 2012;303:R639–50.

62. Woodruff ER, Chun LE, Hinds LR, Spencer RL. Diurnal Corticosterone Presence and Phase Modulate Clock Gene Expression in the Male Rat Prefrontal Cortex. Endocrinology. 2016;157:1522–1534.

63. Saper CB, Loewy AD, Swanson LW, Cowan WM. Direct hypothalamo-autonomic connections. Brain Res. 1976;117:305–312.

64. Lisa M, Marmo E, Wible JH, DiMicco JA. Injection of muscimol into posterior hypothalamus blocks stress-induced tachycardia. Am J Physiol. 1989;257:R246–51.

65. Zhou J-J, Ma H-J, Shao J, Wei Y, Zhang; Xiangjian, Zhang Y, De-;, Li P. Downregulation of Orexin Receptor in Hypothalamic Paraventricular Nucleus Decreases Blood Pressure in Obese Zucker Rats. J Am Heart Assoc. doi:10.1161/JAHA.118.011434.

66. Myers B. Corticolimbic regulation of cardiovascular responses to stress. Physiol Behav. 2017;172:49–59.

67. McKlveen JM, Moloney RD, Scheimann JR, Myers B, Herman JP. ‘Braking’ the Prefrontal Cortex: The Role of Glucocorticoids and Interneurons in Stress Adaptation and Pathology. Biol Psychiatry. 2019. doi:10.1016/J.BIOPSYCH.2019.04.032.

68. Möller-Leimkühler AM. Gender differences in cardiovascular disease and comorbid depression. Dialogues Clin Neurosci. 2007;9:71–83.

69. Tobet SA, Handa RJ, Goldstein JM. Sex-dependent pathophysiology as predictors of comorbidity of major depressive disorder and cardiovascular disease. Pflügers Arch - Eur J Physiol. 2013;465:585–594.

70. Pimple P, Lima BB, Hammadah M, Wilmot K, Ramadan R, Levantsevych O, Sullivan S, Kim JH, Kaseer B, Shah AJ, Ward L, Raggi P, Bremner JD, Hanfelt J, Lewis T, Quyyumi AA, Vaccarino V. Psychological Distress and Subsequent Cardiovascular Events in Individuals With Coronary Artery Disease. J Am Heart Assoc. 2019;8:e011866.

71. Brooks SD, Hileman SM, Chantler PD, Milde SA, Lemaster KA, Frisbee SJ, Shoemaker JK, Jackson DN, Frisbee JC. Protection from vascular dysfunction in female rats with chronic stress and depressive symptoms. Am J Physiol Circ Physiol. 2018;314:H1070–H1084.

72. Hao Z, Jiang X, Sharafeih R, SHEN S, Hand AR, Cone RE, O’Rourke J. Stimulated release of tissue plasminogen activator from artery wall sympathetic nerves: Implications for stress-associated wall damage. Stress. 2005;8:141–149.

73. Brown IAM, Diederich L, Good ME, DeLalio LJ, Murphy SA, Cortese-Krott MM, Hall JL, Le TH, Isakson BE. Vascular Smooth Muscle Remodeling in Conductive and Resistance Arteries in Hypertension. Arterioscler Thromb Vasc Biol. 2018;38:1969–1985.

74. Swanson LW. Brain Maps: Structure of the Rat Brain, Third Ed. Los Angleles, CA, USA: Academic Press; 2004: 1-1–215.

